## Supplementary Materials and Methods for "PlasticEnz: An integrated database and screening tool combining homology and machine learning to identify plastic-degrading enzymes in meta-omics datasets"

1. *Machine Learning Model Selection and Evaluation*

We tested three ML classifiers: a neural network, Random Forest and XGBoost. In all cases, the input features were precomputed protein embeddings and a one-hot encoded feature matrix. Training and test data were loaded from pickle files, and the training set was split (80/20) to create a validation set. A single hidden‐layer feed-forward network (ReLU activation, 128 or 256 units) with dropout (0.3 or 0.5) was trained via Adam (learning rate 1×10⁻³ or 5×10⁻⁴; weight decay 1×10⁻⁴), using weighted BCEWithLogitsLoss. We performed an 80∕20 train/validation split, exhaustively grid-searched hidden size, dropout, and learning rate over up to 100 epochs with early stopping (patience = 10) based on validation loss. The best model was then retrained on the full training set and evaluated on the unseen test set using ROC AUC, accuracy, precision, recall, and F1‐score using scikit-learn’s metric functions (Supplementary Text 1). Everything from the model definition (torch.nn), data loading (torch.utils.data), loss (nn.BCEWithLogitsLoss), optimizer (torch.optim.Adam), and training loops was done using the PyTorch (torch, v.2.2.2) library. We implemented a multi‐label Random Forest classifier using the RandomForestClassifier class from scikit-learn (v. 1.3.2). To handle multiple binary targets simultaneously, we wrapped the base estimator in MultiOutputClassifier. We performed an exhaustive grid search over three hyperparameters—number of trees (n_estimators ∈ {100, 200}), maximum tree depth (max_depth ∈ {None, 5, 10}), and minimum samples per split (min_samples_split ∈ {2, 5})—using ParameterGrid(). The training set was split 80/20 into training and validation subsets via train_test_split, and for each hyperparameter combination we fit on the training fold and computed the mean F1-score across all labels on the validation fold. The combination yielding the highest average F1 was selected, the corresponding Random Forest was retrained on the full training set, and its performance was assessed on the independent test set by computing per-label ROC AUC, precision, recall, F1-score and accuracy with scikit-learn’s metric functions. We trained an XGBoost classifier through the XGBClassifier API (from the xgboost library, v. 2.1.3) with objective binary:logistic and evaluation metric logloss. Again, we used MultiOutputClassifier to extend to multi-label outputs. Hyperparameter tuning explored n_estimators ∈ {100, 200}, max_depth ∈ {3, 5, 7}, and learning_rate ∈ {0.1, 0.05, 0.01} via ParameterGrid(). Using the same 80/20 train/validation split, each candidate was trained on the training fold and evaluated by average F1-score on the validation fold; the best set was then used to retrain on the full training data. Final test‐set evaluation mirrored that of the Random Forest, computing ROC AUC, precision, recall, F1-score, and accuracy for each label with scikit-learn. Final models were saved (neural network as a PyTorch state dictionary, Random forest and XGBoost as pickle files), and detailed metric values, including confidence intervals, are provided in *Table 1*. XGBoost and Neural networks are included in the *PlasticEnz* prediction module which provides prediction scores for each class. For XGBoost, prediction scores correspond to the class probability for each target polymer (PET or PHB) extracted from the second column of predict_proba() (scikit-learn, v. 1.3.2) . For the neural network, prediction scores are computed by applying a sigmoid activation to the model logits, returning class-specific probabilities for each polymer.

2. *Evaluation sets*

To test our ML predictions, we applied our tool to the Illumina whole-genome metagenomics paired-end data from Pinnell et al. (2019) study[^[41]^](https://www.zotero.org/google-docs/?j73IbY) . The study investigated benthic, free-swimming biofilm communities developed on ceramic, PET, and PHA beads over 28‐days during a microcosm experiment at a hypersaline Laguna Madre lagoon (Texas Gulf Coast, USA). The authors reported that while biofilms on PET and ceramic substrates were similar, the PHA-associated biofilms were distinct and characterized by an enrichment of PHB degradation enzymes and overall increase in depolymerase, esterase, lipase and cutinase enzymes. Moreover, a subsequent 15-month exposure study demonstrated visible weight loss of the PHA pellets, indicative of an active PHA-degrading community. Our tool was applied to these datasets to identify and compare plastic-degrading enzyme homologs across exposure conditions. We report results of *PlasticEnz* tested on three groups of samples; bacterial community extracted from PET beads (PET group), from PHA beads (PHB group), from ceramic beads (ceramic) and negative control represented by bacterioplankton extracted from the surrounding seawater (H_2_O). Raw reads were downloaded from European Nucleotide Archive (ENA)[^[42]^](https://www.zotero.org/google-docs/?c85HQp), BioProject accession: PRJEB15404. All reads were quality-filtered with Fastp[^[43]^](https://www.zotero.org/google-docs/?wppPho) (default settings) and assembled into contigs using Megahit (default settings)[^[44]^](https://www.zotero.org/google-docs/?5reJfY). The resulting contigs served as inputs for the *PlasticEnz* analysis pipeline. To account for differences in sequencing depth among samples, counts of identified PHB/PET depolymerase homologs were normalized against the total number of predicted proteins per sample, and expressed as hits per million proteins for heatmaps and bar charts. To compare similarities between the highest-scoring *PlasticEnz* predictions for PHB and PET degradation and confirmed depolymerases from our database, we extracted the top 10 predictions from each model (PHB-default, PHB-sensitive, and PET-sensitive). For PET biofilms, only PET-sensitive predictions were available. These predicted sequences, derived from their respective biofilms, were combined with non-redundant SEED reference enzymes from the *PlasticEnz* database (clustered at 90% identity to prevent redundancy using CDHIT2[^[38]^](https://www.zotero.org/google-docs/?spjwz5)). Combined sequences were aligned using MUSCLE (default settings)[^[45]^](https://www.zotero.org/google-docs/?9RCExa), and poorly aligned regions were removed using trimAl (v1.4.1) (-automated1 option)[^[46]^](https://www.zotero.org/google-docs/?odJFuV). Pairwise evolutionary distances were calculated with the LG substitution model[^[47]^](https://www.zotero.org/google-docs/?3k4eSR) implemented by the phangorn package (v2.12.1) in R. The resulting distance matrix was then scaled with min-max normalization to generate similarity scores (ranging from 0 to 1) and visualized as heatmaps using the pheatmap R package (v1.0.12).

To test our custom-designed HMM motifs, we applied the *PlasticEnz* tool to publicly available metagenomic datasets from two contrasting environments. The first environment consisted of thermophilic bacterial communities from pristine acidic hot springs in the Kamchatka Peninsula, Russia, an area characterized by extreme conditions and geographic remoteness, making plastic contamination unlikely. Sediment samples were collected from the Mutnovsky volcano hot spring at a depth of one meter, an elevation of approximately 2,000 meters, and a temperature of around 70°C (NCBI BioProject accession PRJNA419239; samples SRR6311220, SRR6311221, SRR6311224). The second set of environments included soil microbiomes collected from urban dumping sites in Iran and India with documented long-term plastic contamination. Specifically, this data set included soil samples from the Sewapura landfill in Jaipur, India ("Sewapura"), which were collected at approximately 0.6 meters depth, with plastic waste accumulation occurring over several years (BioProject accession PRJNA1077790; sample SRR28006989)[^[48]^](https://www.zotero.org/google-docs/?cFhoZ9) and agricultural soils from Varamin and Ghaleh-No village in Tehran province, Iran ("Varamin") which were exposed to plastic mulch and municipal wastewater contamination for approximately 35 years (BioProject accession PRJNA924045; sample SRR23085642)[^[49]^](https://www.zotero.org/google-docs/?4mT9gT). Lastly, we analysed soil samples from the Ghazipur landfill site in Delhi, India ("Ghazipur"), which receives approximately 2,200 metric tonnes of waste daily (BioProject accession PRJNA388130; sample SRR5613220). Raw sequencing data (paired-end or single-end short reads) were downloaded from ENA and processed using the same pipeline as Laguna Madre microcosm samples. To investigate sequence-level similarity between *PlasticEnz*-predicted homologs and known enzyme SEED sequences from the *PlasticEnz* seed database, Principal Coordinates Analysis (PCoA) was performed. Putative PETase sequences from contaminated sites (Varamin, Sewapura, Ghazipur) with high prediction score (default model) for PET or with a HMMER bitscore > 80 were combined with corresponding SEED sequences. Combined protein fasta were aligned using Clustal Omega (version 1.2.4)[^[50]^](https://www.zotero.org/google-docs/?kXs5zF), and pairwise Fitch distances[^[51]^](https://www.zotero.org/google-docs/?Jvg98O) between sequences were calculated with *dist.alignment* (R-package: seqinr)[^[52]^](https://www.zotero.org/google-docs/?uHCrPK). PCoA ordination was performed with the *pcoa* (R-package: *ape*[^[53]^](https://www.zotero.org/google-docs/?Gntypm)). Phylogenetic trees of previous Clustal Omega alignments were trimmed with TrimAl (gap threshold of 0.5) and constructed with FastTree[^[54]^](https://www.zotero.org/google-docs/?vSSKKM) (LG model).

All data analysis was performed in R (v. 4.2) and python (v. 3.11.11). All bar charts and heatmaps were created using ggplot2, ggpubr, and patchwork packages
