## Supplementary Text 1 for "PlasticEnz: An integrated database and screening tool combining homology and machine learning to identify plastic-degrading enzymes in meta-omics datasets"

### Evaluation Metrics

#### Precision

$$\text{Precision} = \frac{TP}{TP + FP}$$

#### Recall

$$\text{Recall} = \frac{TP}{TP + FN}$$

#### F1-score

$$\text{F1} = 2 \cdot \frac{\text{Precision} \cdot \text{Recall}}{\text{Precision} + \text{Recall}} = \frac{2TP}{2TP + FP + FN}$$
